# A divergent betacoronavirus with a functional furin cleavage site in South American bats

**DOI:** 10.1101/2025.10.24.684489

**Authors:** Kosuke Takada, Nicholas Yamahoki, Jonathon C. O. Mifsud, Itsuki Anzai, Tadashi Maemura, Francisco Borges Costa, Eric Takashi Kamakura de Carvalho Mesquita, Mateus de Souza Ribeiro Mioni, Tiago JS Lopes, Yoshihiro Kawaoka, Jane Megid, Edward C. Holmes, Tokiko Watanabe

## Abstract

Bats are natural reservoirs for a wide range of RNA viruses. Members of the genus *Betacoronavirus*, including Severe Acute Respiratory Syndrome virus 2 (SARS-CoV-2) and Middle East Respiratory Syndrome virus (MERS-CoV), have attracted particular attention due to their recent zoonotic emergence. However, much of the known diversity of betacoronaviruses is based on data from Asia, Africa, and the Middle East, with limited genomic information available from the Americas. Herein, we report the complete genome of a novel bat betacoronavirus identified from a *Pteronotus parnellii* bat sampled in Brazil. Phylogenetic analysis revealed that this virus is sufficiently distinct from the five recognized *Betacoronavirus* subgenera to represent a new subgenus. Of note, the spike protein of this novel bat coronavirus possesses a functional furin cleavage site at the S1/S2 junction with a unique amino acid sequence motif (RDAR) that differs from that found in SARS-CoV-2 (RRAR) by only one amino acid. Comparative structural analysis identified other betacoronaviruses in bats with furin cleavage sites at the S1/S2 junction, suggesting that this region is a structurally permissive “hotspot” for cleavage site incorporation. Our study provides a broader understanding of the phylogenetic and functional diversity of bat coronaviruses as well as their zoonotic potential.

## Introduction

Bats (order *Chiroptera*) are among the most diverse groups of mammals^1,2^ and similarly harbor a diverse array of RNA viruses, including coronaviruses and paramyxoviruses with zoonotic potential^3–5^. Various members of the genera *Alphacoronavirus* and *Betacoronavirus* have been identified in bats, which are considered important natural hosts for these viruses^3,5,6^, and the betacoronaviruses SARS-CoV, SARS-CoV-2, and MERS-CoV recently emerging in humans^7,8^. The high diversity of viruses in bats therefore positions them as a key taxonomic group for zoonotic disease surveillance^9^. Although metagenomic-based surveys have identified many bat-associated viruses, the viral sequences obtained often only comprised the RNA-dependent RNA polymerase (RdRp) used in RNA replication, with limited coverage of other viral proteins (e.g., surface proteins) that typically determine host range, tissue tropism, and pathogenicity.

Although novel RNA viruses continue to be identified in bats, especially in China and South-East Asia^5,6^, our understanding of the phylogenetic diversity of bat-associated RNA viruses is also limited a bias in sampling toward particular geographic localities. For example, our current knowledge of the diversity of betacoronaviruses is largely based on sampling from Asia, Africa, and the Middle East^10,11^. In marked contrast, although South America is classified as a global biodiversity hotspot^12^, relatively little is known about the RNA viruses that circulate in bats in this large geographic region.

Coronaviruses (CoVs) are characterized by the presence of a spike (S) surface glycoprotein that binds to receptors on the cell surface to initiate infection. Some CoV S proteins are also characterized by the presence of a furin cleavage site (FCS) at the S1/S2 junction, in which the S1 subunit is responsible for receptor binding and the S2 subunit mediates membrane fusion^13^. The FCS facilitates protease cleavage within the host cell, contributing to enhanced infectivity and/or pathogenicity^14–17^. SARS-CoV-2 has an FCS at the S1/S2 junction. While this feature is not found in closely related bat-associated sarbecoviruses^18^, a recent study identified a furin-like cleavage site in bat betacoronaviruses basal to the sarbecovirus lineage^19^, indicating that similar motifs can arise naturally in bats. However, the diversity, evolution, and functional significance of the FCS in bat betacoronaviruses has yet to be systematically investigated. Here, we describe a novel betacoronavirus with a functional FCS that was found in Brazil, a region where surveillance has been limited.

## Results

### Bat sampling in Brazil and sequencing

We captured diverse bat species from three locations in Brazil (Riachão [Maranhão state], Botucatu [São Paulo], and Arari [Maranhão]) (Fig. 1A and Supplementary Table S1), from which we sampled intestinal tissues. Samples were obtained from multiple species of microbats (members of the families *Phyllostomidae* and *Mormoopidae*), including *Anoura caudifer*, *Anoura geoffroyi*, *Carollia perspicillata*, *Desmodus rotundus*, *Glossophaga soricina*, *Pteronotus parnellii*, and *Rhinophylla alethina*, which were identified based on morphological characteristics (Fig. 1A and Supplementary Table S1).

**Figure 1.**
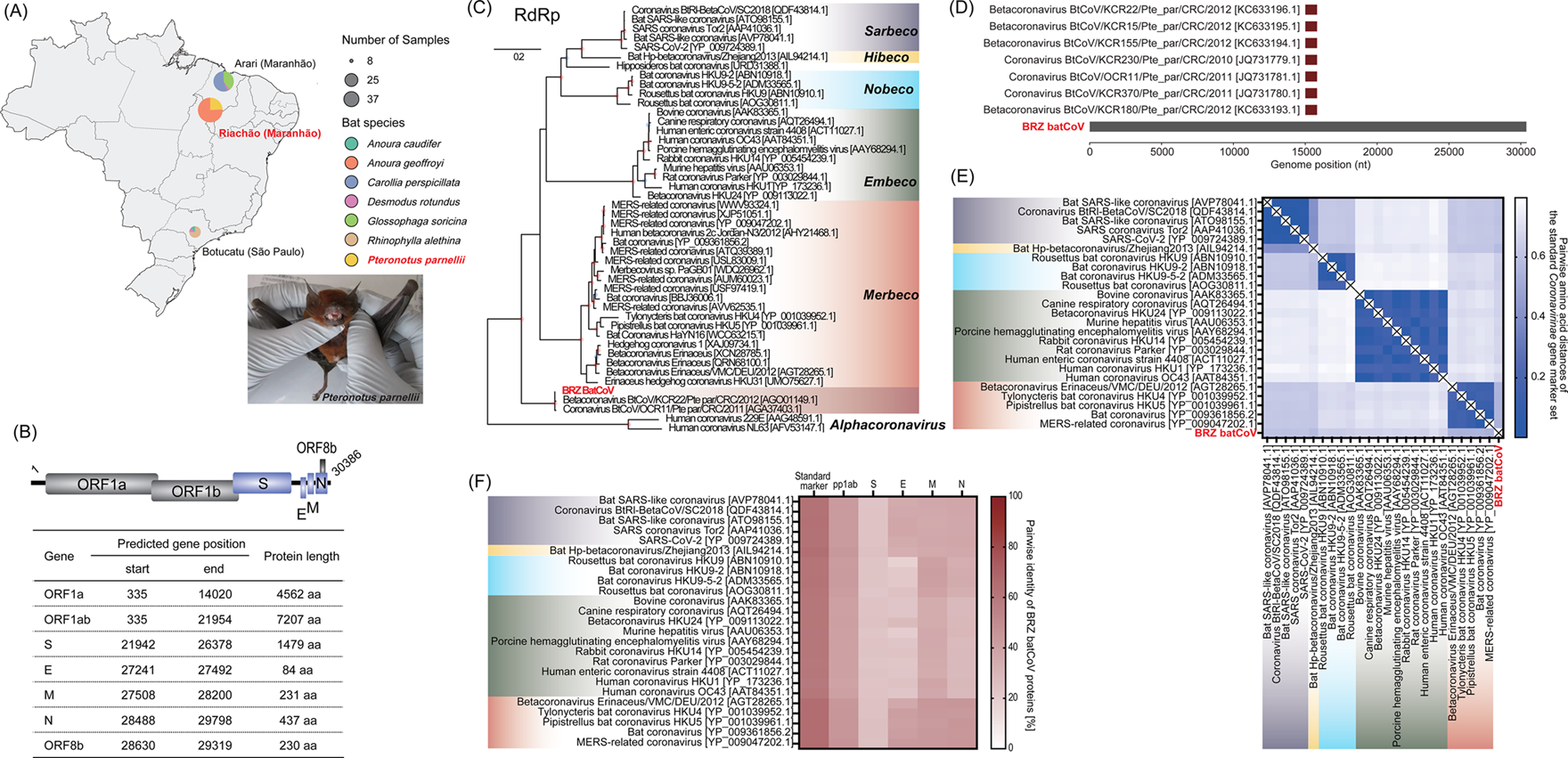
Sampling context, genomic features, and evolutionary relationships of the novel betacoronavirus. (A) Map showing the composition and number of bat samples collected at three locations in Brazil: Riachão (Maranhão), Botucatu (São Paulo), and Arari (Maranhão). The pie charts represent bat species composition, with chart size indicating the number of individual samples collected at each location. Geographic locations where samples with Coronavirus (CoV) genomes were collected and bat species are indicated in red. Representative photograph of the bat species (*Pteronotus parnellii*), in which CoV genomes were detected, is shown. (B) Schematic overview of newly identified viruses belonging to the genus *Betacoronavirus* with corresponding sample information. (C) Maximum likelihood phylogenetic tree inferred from amino acid sequences of the (RdRp (non-structural protein 12), including the virus identified in this study, representative viruses, and viruses showing the greatest similarity in BLAST, including those matching partial sequences. Branch lengths represent the number of substitutions per site. Red and blue circles at internal nodes indicate bootstrap values ≥ 90% and ≥ 80%, respectively. Each subgenus within the genus *Betacoronavirus* is represented by a different color. The red characters denote the CoV identified in this study (BRZ batCoV). (D) Nucleotide alignment between the newly identified virus and the most closely related partial RdRp sequences. (E) Pairwise genetic distances of the standard *Coronavirinae* gene marker set (nsp5, nsp12, nsp13, nsp14, nsp15, and nsp16 protein) in representative betacoronaviruses. The number of amino acid substitutions per site from between sequences is shown. (F) Pairwise amino acid sequence identity between the newly identified virus and representative betacoronaviruses.

We generated 16 metatranscriptomic libraries from pooled intestinal RNA of bats representing different species and collection regions. Each library yielded approximately 40 million paired-end reads on the Illumina NovaSeq 6000 platform. We assembled the quality-filtered reads into contigs and screened them against the GenomeSync database (http://genomesync.org). Contigs with top hits to viral genomes were considered virus-like and manually curated, focusing on identifying CoV-like sequences.

### Genomic features and evolutionary relationships of the novel betacoronavirus

Through metagenomic sequencing we identified a partial CoV-like genome from a library of Parnell’s mustached bat (*Pteronotus parnellii*) collected at Place Riachão, Maranhão state, Brazil. Because the whole genome may not have been identified due to the mixing of multiple samples, we individually sequenced the samples in the library in which CoV-like sequences were found. This led to the identification of a 30,386-nucleotide CoV genome, tentatively named BRZ batCoV, that shared 66.99% amino acid identity with the ORF1ab polyprotein of *Betacoronavirus* Erinaceus/VMC/DEU/2012 [GenBank ID: YP_009513008.1], a member of the *Merbecovirus* subgenus. Gene annotation of the BRZ batCoV genome was performed based on sequence similarity to annotated reference genomes of merbecoviruses. This analysis identified two large open reading frames [ORF1a (encoding the polyprotein pp1a) and ORF1ab (encoding the polyprotein pp1ab)], as well as genes encoding the S, envelope (E), membrane (M), nucleocapsid (N), and ORF8 proteins (Fig. 1B and Supplementary Table S2-S3).

To determine the evolutionary relationships between BRZ batCoV and other betacoronaviruses we inferred phylogenetic trees based on the amino acid sequence of the nsp12 protein that contains the RdRp, using alphacoronaviruses as an outgroup. This yielded a CoV phylogeny in which each of the five recognized subgenera (*Sarbecovirus*, *Hibecovirus*, *Nobecovirus*, *Merbecovirus*, and *Embecovirus*) formed a distinct cluster with strong bootstrap support (Fig. 1C). In contrast, BRZ batCoV formed a distinct and well supported cluster (Fig. 1C) with two bat CoVs [BtCoV/OCR11/Pte par/CRC/2011 (GenBank ID: AGA37403.1) and BtCoV/KCR22/Pte par/CRC/2012 (GenBank ID: AGO01149.1)] sampled from the same bat species in Costa Rica, but for which only partial RdRp sequences (816-nucleotide)^20^ were available (Fig. 1D).

To better evaluate the genetic distinctiveness of BRZ batCoV we computed amino acid genetic distances in the standard *Coronavirinae* gene marker set (nsp5, nsp12, nsp13, nsp14, nsp15, and nsp16 protein). Intra-subgenus genetic distances ranged from 0.002–0.201 within each of the five betacoronavirus subgenera, and from 0.400–0.796 substitutions/site between the subgenera (Fig. 1E and Supplementary Table S4). The genetic distance between BRZ batCoV and other subgenera ranged from 0.541–0.704 substitutions/site (Fig. 1E and Supplementary Table S4), with similar values in the S, E, M, and N proteins (Supplementary Fig. S1 and Supplementary Table S5-S8). Hence, BRZ batCoV is sufficiently genetically distinct from known betacoronaviruses to represent a novel subgenus.

To more precisely document the evolutionary position of BRZ batCoV, we constructed individual amino acid alignments of the ORF1ab, S, E, M, and N proteins with representative betacoronaviruses and conducted genetic distance computations and phylogenetic analyses (Fig. 1F, Fig. 2 and Supplementary Table S9). The standard *Coronavirinae* marker set of BRZ batCoV showed the highest amino acid sequence identity (65.8%) to Middle East respiratory syndrome-related CoV [YP_009047202.1], which was also the most closely related virus with respect to the polyprotein1ab (45.7%) and the E protein (43.4%). For the M protein, the highest identity was 44.8% to Betacoronavirus Erinaceus/VMC/DEU/2012 [AGX27817.1], which was also the most closely related virus with respect to the N protein (45.4%). In contrast, the S protein showed generally low identity to representative viruses, with the highest being 25.7% to Pipistrellus bat CoV HKU5 [YP_001039962.1]. Taken together, these data confirm that BRZ batCoV is sufficiently genetically distinct to represent a novel subgenus of betacoronaviruses, with particularly pronounced differences observed in the S protein. The distinctiveness of BRZ batCoV was also apparent from phylogenetic analyses of each protein^21^, although it generally shared common ancestry with members of *Merbecovirus* subgenus (Fig. 2).

**Figure 2.**
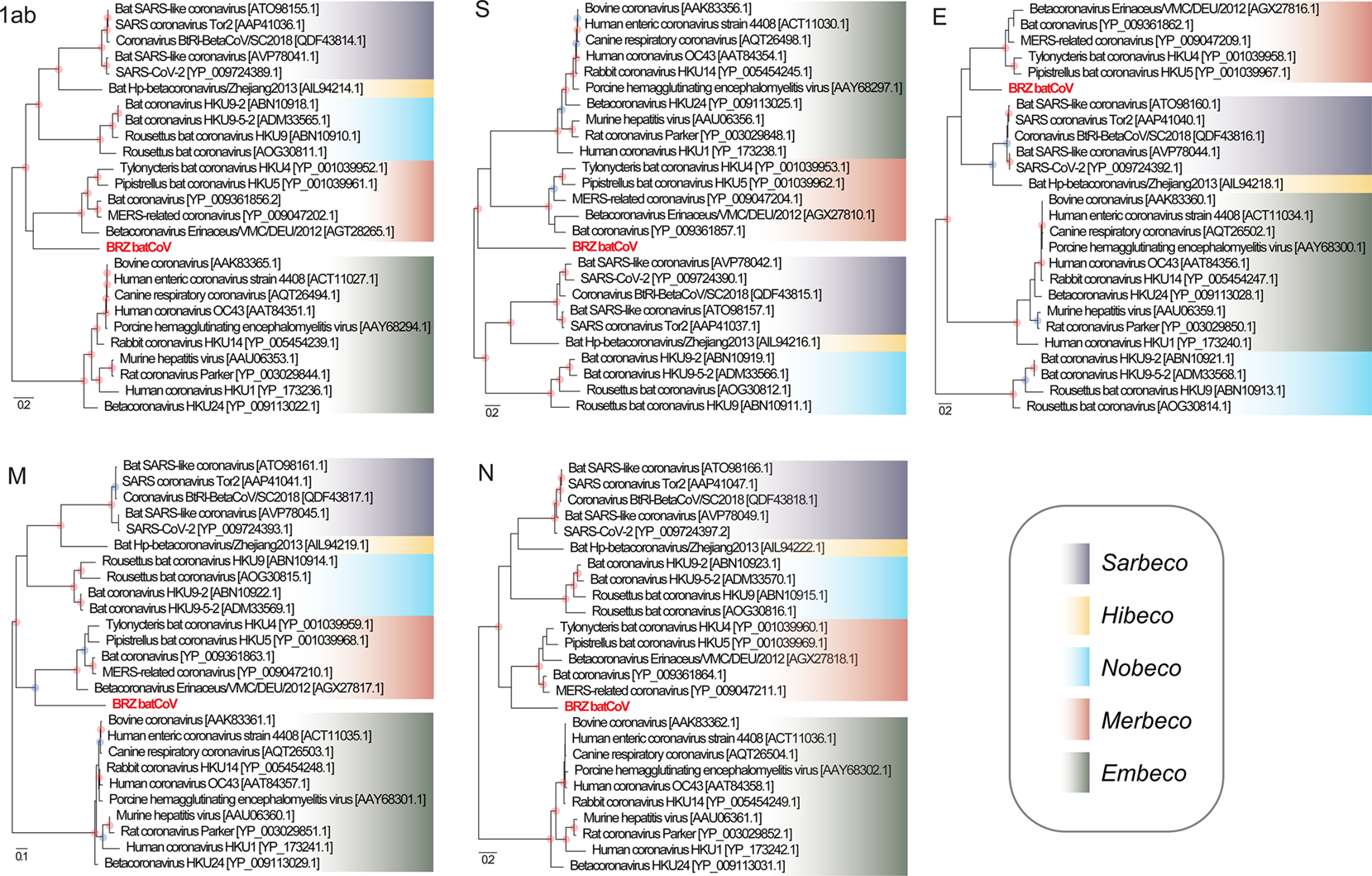
Individual gene phylogenies of the betacoronavirus identified from *Pteronotus parnellii* bats sampled in Brazil. Maximum likelihood phylogenetic trees of amino acid sequences of the ORF1ab, S, E, M and N proteins, including the virus identified in this study and representative viruses. Branch lengths represent the number of amino acid substitutions per site. Red and blue circles at internal nodes indicate bootstrap values ≥ 90% and ≥ 80%, respectively. Each subgenus of betacoronaviruses is represented by a different color. Red characters indicate BRZ batCoV, which was identified in this study.

### Structural prediction of the BRZ batCoV S protein

Since the CoV S glycoprotein is a key determinant of host range and pathogenicity, we next investigated the properties of the S protein of BRZ batCoV. We predicted the tertiary protein structures for the BRZ batCoV as both a monomer and a homotrimer by using the AlphaFold3^22^. Despite a relatively low average predicted local distance difference test (pLDDT) score of 58, we observed broad-scale structural similarity with the S glycoprotein structures of other betacoronaviruses (Fig. 3A and 3B). For example, when aligned with Hedgehog CoV 1 (9JMG) several structural blocks were moderately conserved, including those that spanned the S1 C terminal domain and into the beginning of the S2 subunit (unpruned atom RMSD = 4.93 Å, residues 809–1,018), with the remaining S2 subunit displaying greater structural similarity (unpruned atom RMSD = 2.80 Å, residues 1,025–1,352) (Supplementary Fig. S2A). These regions also corresponded to the most confidently predicted regions in the BRZ batCoV structure based on predicted aligned error (PAE) and pLDDT scores (Supplementary Fig. S2B and S2C).

**Figure 3.**
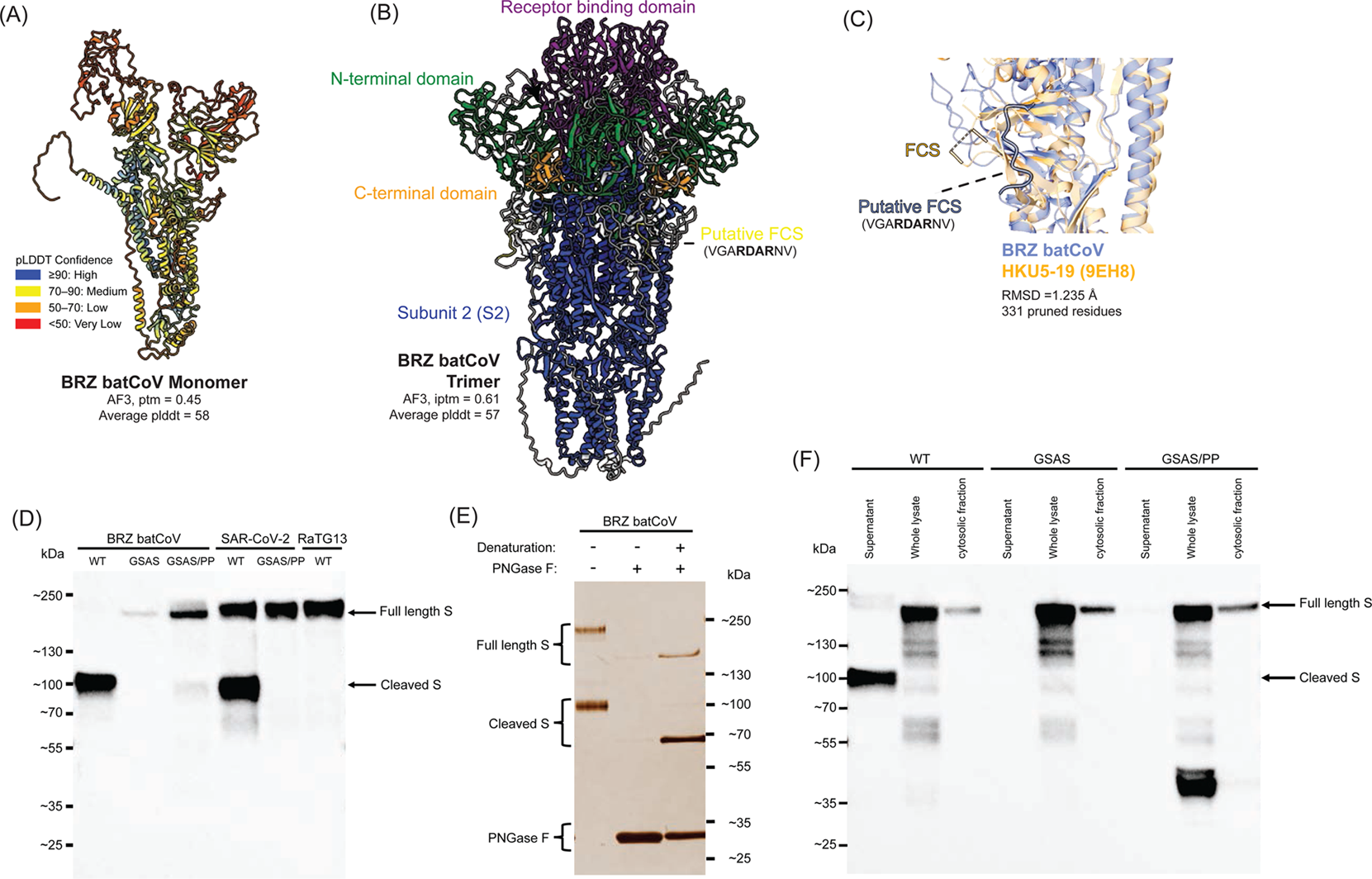
Characterization of the S protein of BRZ batCoV. (A) Predicted protein structure of the BRZ batCoV S glycoprotein monomer color-coded by pLDDT confidence scores as shown in the key. (B) Predicted BRZ batCoV S glycoprotein trimer with select domains colored: receptor-binding domain, purple; N-terminal domain, green; C-terminal domain, orange; and subunit 2, blue. (C) Structural superposition of the predicted BRZ batCoV S monomer and experimentally determined HKU5-19 structure (PDB 9EH8) using Matchmaker focused on the S1/S2 junction, with the potential furin cleavage site (FCS) annotated. The FCS in 9EH8 (residues 751-758) was not modeled, but the modeled region surrounding this (residues -751 and 765-) was highlighted to show its approximate position. (D) Western blotting of the S protein expressed from plasmids in Expi293F cells. S proteins from BRZ batCoV (wild-type, GSAS mutant, and GSAS/PP mutant), SARS-CoV-2 (wild-type and GSAS/PP mutant), and bat RaTG13 (wild-type) were analyzed after purification by nickel affinity chromatography. All constructs were engineered as soluble ectodomains lacking transmembrane and cytoplasmic domains, with C-terminal His-tags for detection with an anti-His tag antibody. GSAS mutations replaced the FCS motif (RDAR in BRZ batCoV; RRAR in SARS-CoV-2) with a non-cleavable sequence. PP denotes double proline mutations introduced to stabilize the prefusion trimer. Arrows indicate full-length S and cleaved S2 fragments. (E) Treatment of purified BRZ batCoV S with peptide-N-glycosidase F (PNGase F) under non-denaturing and denaturing conditions. PNGase F removes N-linked glycans, allowing visualization of the deglycosylated protein core and confirmation of predicted molecular weights (full-length S approx. 162 kDa; S2 fragment approx. 69 kDa). (F) Western blotting comparing intracellular and extracellular localization of BRZ batCoV S protein cleavage. Expi293F cells expressing wild-type or mutant (GSAS or GSAS/PP) BRZ batCoV S were fractionated into cytosolic fractions and culture supernatants. Whole cell lysates are shown for reference. His-tagged S proteins were detected by using an anti-His tag antibody.

BRZ batCoV was particularly notable in that it possesses a putative furin cleavage site at amino acid position 831, located near the S1/S2 junction of the S protein (Supplementary Table S10). Strikingly, the BRZ batCoV sequence contains a cleavage site motif – RDAR – that differs from that of MERS-CoV (RSVR), but which is only one amino acid substitution away from that seen in SARS-CoV-2 (RRAR) (Supplementary Table S11). As the FCS in BRZ batCoV is located upstream of where FCSs are typically seen in the S1/S2 junction of other betacoronaviruses, we examined whether it occupied a similar position structurally. The structure predicted by AlphaFold3 was moderately confident in the FCS prediction in both the BRZ batCoV monomer and trimer models (average pLDDT for residues 825-833 = 78.89 and 75.69, respectively). Specifically, the BRZ batCoV FCS appears as an exposed loop in the trimer model, whereas in the monomer model a short β-sheet was predicted within this loop. Hence, the FCS appears structurally located in the same region as that for other betacoronaviruses (Fig. 3C), albeit with a slight positional offset reflected in the amino acid alignment.

### Functional characterization of the putative FCS

To experimentally validate the predicted FCS, we expressed recombinant BRZ batCoV S protein in Expi293F cells. Protein samples were purified by nickel affinity chromatography and then subjected to sodium dodecyl sulfate-polyacrylamide gel electrophoresis (SDS-PAGE) and analyzed by Western blotting using an anti-His tag antibody. A band corresponding to the full-length BRZ batCoV S was observed as was a lower molecular weight-protein band (Fig. 3D). Removal of the predicted FCS prevented cleavage of the BRZ batCoV S, as indicated by the reduced intensity of the lower molecular weight band. As expected, the same phenomenon was observed for SARS-CoV-2 S, which has an FCS at the S1/S2 junction. In contrast, the S protein of bat sarbecovirus RaTG13, which lacks an FCS between its S1 and S2 subunits, was resistant to protease cleavage. These results demonstrate that proteolysis of BRZ batCoV S is due to the presence of a functional furin cleavage site.

Based on the *in silico* analysis, the full-length BRZ betaCoV S has a molecular weight of approximately 162 kDa, and cleavage at the predicted FCS should generate an S2 fragment with a molecular weight of about 69 kDa. To confirm this, the purified BRZ batCoV S was treated with peptide-N-glycosidase F (PNGase F) and the migration of the deglycosylated S was then evaluated by SDS-PAGE. The result was consistent with the *in silico* prediction (Fig. 3E). We also investigated the site of S cleavage. Expi293F cell pellets were lysed and analyzed by SDS-PAGE and Western blotting. The result demonstrated that cleavage of the BRZ batCoV S protein occurred predominantly after its secretion into the culture medium, with no detectable cleavage in the cytosolic fraction (Fig. 3F). Interestingly, mutating the FCS motif from RDAR to GSAS in BRZ batCoV S reduced the level of secretion, although the underlying reason is currently unclear.

The S proteins were further purified by size-exclusion chromatography (Supplementary Fig. S3A-C). From 120-mL cultures of Expi293F cells, 298.9 μg/mL of SARS-CoV-2 S(GSAS/PP) and 237.0 μg/mL of RaTG13 S(WT) were obtained (Supplementary Fig. S3D). The yield of BRZ batCoV S(WT) was 4.3- and 3.5-fold lower than that of SARS-CoV- 2 S and RaTG13 S, respectively. These results suggest that BRZ batCoV S might be less efficiently expressed in human cells compared to SARS-CoV-2 S and RaTG13 S.

### Phylogenetic distribution of FCSs in betacoronaviruses

We next investigated whether other bat-derived betacoronaviruses possess a putative FCS similar to that identified in BRZ CoV. We used ProP to predict putative FCSs near the S1/S2 junction (i.e., the amino acid region spanning S protein residues 599–900) in betacoronaviruses with annotated host species. Putative FCSs were detected in one of 169 sequences of the subgenus *Sarbecovirus* [Bat SARS-like CoV Khosta-1 (QVN46559.1)], two of two sequences from the subgenus *Hibecovirus*, and 11 of 26 sequences from the subgenus *Merbecovirus* (Fig. 4A and Supplementary Table S11). These results are consistent with previous reports showing that FCSs have emerged multiple times in coronavirus evolution^23^.

**Figure 4.**
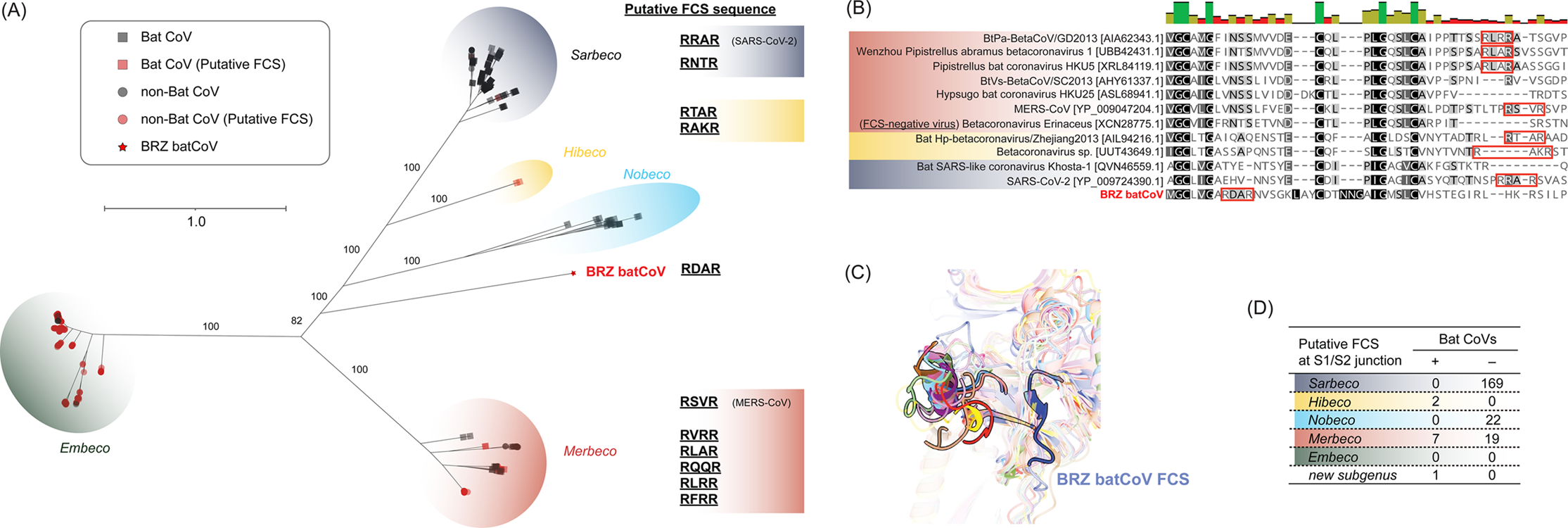
Putative furin cleavage sites in bat-derived betacoronaviruses and their structural locations. (A) Detection of putative furin cleavage sites in bat-derived betacoronaviruses. Full-length S protein sequences were analyzed using ProP to predict potential FCSs near the S1/S2 junction based on each amino acid position (i.e., the amino acid region spanning residues 599–900). The presence or absence of predicted FCSs is summarized by subgenus, highlighting distinctions between bat-derived and non-bat-derived CoVs. (B-C) Structural and sequence comparison of the S1/S2 junction in SARS-CoV-2. (B) Pairwise amino acid alignment between the sequences underlying the structures in (C) with the FCS annotated. (C) Superposition of predicted betacoronavirus S proteins with the predicted FCSs highlighted. The viruses used for superposition are shown in Supplementary Table S11. (D) Number of bat CoVs with an FCS at a similar S1/S2 junction in SARS-CoV-2, shown by subgenus.

Following the amino acid alignment of the S proteins of bat-derived viruses, some of the putative FCSs at the S1/S2 junction did not align to the same amino acid position (Fig. 4B), instead falling at atypical positions such as near the N-terminal region of S1 or within S2. As with BRZ batCoV, we assessed their structural location by using AlphaFold3-predicted structures. Across the five known subgenera, nine of 15 other betacoronaviruses for which we predicted the structures of their S proteins (including one excluded from structural prediction but inferred from sequence alignment; see Supplementary Table S11) possessed FCSs that were structurally located in the same region at the S1/S2 junction, similar to that of BRZ batCoV, and consistent with those in SARS-CoV-2 and MERS-CoV (Fig. 4C and 4D and Supplementary Table S11). Therefore, the FCSs of these bat-derived CoVs likely undergo cleavage that would result in two S protein subunits, as experimentally demonstrated in SARS-CoV-2 and BRZ batCoV (Fig. 4C and 3D). These results suggest that FCS acquisition events have occurred independently multiple times in bat betacoronaviruses, and that there may be structural “hotspots” for FCS acquisition.

## Discussion

We identified a full-length genome of a novel bat CoV (BRZ batCoV) from a *Pteronotus parnellii* bat sampled in Brazil that is phylogenetically distinct from known betacoronaviruses. This lineage originates from an under-sampled geographic region, thereby expanding the landscape of bat CoV diversity. The phylogenetic distinctiveness of BRZ batCoV suggests that it may represent a new subgenus of betacoronaviruses. Although partial RdRp sequences from phylogenetically related viruses have been reported previously from the same bat species in Costa Rica^20^, the lack of corresponding structural and accessory gene sequences hindered a complete understanding of their genome organization and phylogenetic position. By providing a complete gene set and demonstrating consistent phylogenetic placement across the genome, we were able to address this gap. Of note, BRZ batCoV possesses a furin cleavage site with the sequence motif RDAR, which has not previously been reported in coronaviruses and is only one amino acid different from that seen in SARS-CoV-2 (RRAR). Given the importance of the furin cleavage site in determining host range, infectivity, and cross-species transmission^24,25^, this finding provides important insights into the evolutionary potential and zoonotic risk of BRZ batCoV. More broadly, these results imply that other furin cleavage sites, such as that in SARS-CoV-2, may be acquired in bats by recombination or insertion mutations, further highlighting the role of bats as potential reservoirs of genetic innovations relevant to zoonotic emergence.

Furin cleavage sites have been identified in the surface proteins of other RNA viruses. For example, in highly pathogenic avian influenza viruses (*Orthomyxoviridae*), stem-loop RNA secondary structures might facilitate the insertion of a polybasic cleavage site in the hemagglutinin gene^26,27^, allowing cleavage by ubiquitous cellular proteases such as furin, and in turn enabling replication in tissues beyond the respiratory and intestinal tract, directly contributing to pathogenicity^28–30^. Similarly, in avian orthoavulaviruses 1 (formerly Newcastle disease virus, *Paramyxoviridae*), the polybasic furin-recognition motif at the fusion protein cleavage site is a well-established determinant of virulence, distinguishing highly pathogenic from low-pathogenic strains^31,32^. Ebola virus (*Filoviridae*) glycoproteins also harbor FCSs, although these are not strictly required for replication or virulence in nonhuman primates^33–35^. These examples underscore FCS acquisition as an evolutionary adaptation that independently arises in diverse RNA virus families, often associated with enhanced pathogenicity.

Several molecular evolutionary mechanisms have been proposed for the acquisition of the FCS in coronavirus S proteins. First, short insertion mutations could result in the formation of novel polybasic motifs, as exemplified by the PRRA 12-nt insertion at the S1/S2 junction in SARS-CoV-2^23,36,37^. Second, recombination-mediated introduction can also lead to motif acquisition, such that FCS-like sequences can be acquired through homologous recombination between different CoVs or with host-derived sequences^38–40^. Although CoVs, such as SARS-CoV-2, harbor extensive RNA secondary structures across their genomes^41,42^ that could facilitate mutation and recombination in a specific region^43^, our results suggest that FCS acquisition is impacted by protein-level constraints, with the S1/S2 junction representing a structurally permissive “hotspot” for stable incorporation of cleavage motifs. Moreover, although it has been suggested that the acquisition of the FCS in SARS-CoV-2 may have occurred in an “intermediate host” rather than bats^18^, our study suggests a viable route for it to have emerged within bat populations through recombination. Indeed, bats harbor a highly diverse range of CoVs, providing an evolutionary environment favorable for frequent recombination. Nevertheless, the precise molecular mechanisms underlying the acquisition of FCSs remain poorly understood, underscoring the need for further investigation to elucidate the determinants and constraints shaping this important adaptive trait.

Our study has some limitations. A previous study demonstrated that furin-cleavage at the S1/S2 junction destabilizes the SARS-CoV-2 S protein, thereby promoting the open conformation that exposes the receptor-binding domain and enables high-affinity ACE2 interaction^44^. We did not examine the receptor-binding properties and infectivity of the virus, which restricts the extent to which its zoonotic risk can be evaluated. Hence, although the presence of an FCS is clearly significant, any discussion of the zoonotic potential of this virus should be limited. Future investigations are required to elucidate how the FCS in bat CoVs contributes to host range expansion and pathogenicity through molecular functional validation using authentic viruses and/or pseudoviruses.

Overall, this study underscores the importance of investigating RNA viruses in South American bats as a key step toward enhancing our understanding of the phylogenetic and functional diversity of RNA viruses. Our findings highlight the importance of elucidating FCS acquisition events as a central molecular mechanism that shapes the pathogenicity and host range of these viruses.

## Materials and methods

### Sample collection

We captured bat species from three locations in Brazil (Arari and Riachão in Maranhão state and Botucatu in São Paulo state) between May and August 2019, and collected intestinal tissue samples from 70 bats (Supplementary Table S1). Seven different bat species were collected, identified based on their morphological characteristics: *Anoura caudifer* (n = 1), *Anoura geoffroyi* (n = 28), *Carollia perspicillata* (n = 14), *Desmodus rotundus* (n = 2), *Glossophaga soricina* (n = 10), *Pteronotus parnellii* (n = 9), and *Rhinophylla alethina* (n = 6). All tissue samples were stored in RNAlater and then kept at -80°C until use.

### Total RNA metatranscriptomic sequencing

Total RNA was extracted and purified using the RNeasy Mini Kit (QIAGEN). Based on the bat species identified primarily by morphological criteria and collection region, the extracted RNA was pooled into 16 RNA pools, with each pool containing 1–11 samples of the same type (Supplementary Table S1). The preparation of sequencing libraries and rRNA depletion (host and bacterial) was conducted using the TruSeq Stranded Total RNA Library Prep Kit with Ribo-Zero Plus. Total RNA metatranscriptomic sequencing was performed on a NovaSeq 6000 platform with 100 bp paired-end reads at Macrogen Japan Corp. (Tokyo, Japan), generating approximately 40 million reads per library.

### Genome assembly and annotation

Raw reads were obtained from the 16 pools and were adaptor- and quality-trimmed with fastp^45^. The processed reads were then *de novo* assembled using SPAdes v3.15.2^46^ using the default settings. The resulting contigs were queried via blastn against viral genome sequences in the GenomeSync database (http://genomesync.org). The query sequences with the best hits to viral genome sequences were regarded as virus-like contigs and analyzed in detail manually.

To annotate the viral genome, tblastn searches were conducted against the assembled genome sequence using protein sequences from the closely related MERS-CoV as queries. The following proteins were used: 1ab polyprotein (GenBank: YP_009047202.1), spike protein (GenBank: YP_009047204.1), envelope protein (GenBank: YP_009047209.1), membrane protein (GenBank: YP_009047210.1), nucleoprotein (GenBank: YP_009047211.1), NS3 (GenBank: YP_009047205.1), NS4A (GenBank: YP_009047206.1), NS4B (GenBank: YP_009047207.1), NS5 (GenBank: YP_009047208.1), and ORF8b (GenBank: YP_009047212.1). Searches were performed using an e-value threshold of 1e — 5. The resulting sequence alignments informed manual curation of gene annotations. In parallel, putative open reading frames (ORFs) within the verified genome sequences were predicted using Geneious Prime (version 2025.0.3) as detailed in Supplementary Table S2. To estimate the putative mature protein sequences, we used SignalP 6.0 by integrating predicted cleavage sites between viral nonstructural proteins with sequence alignment results of known nonstructural proteins from representative betacoronaviruses (Supplementary Table S3).

### Estimates of evolutionary divergence between betacoronaviruses

We estimated the number of amino acid substitutions per site between different betacoronaviruses, with the resulting pairwise distances shown in Figure 2E and Supplementary Fig. S3. Pairwise distances with standard errors are summarized in Supplementary Tables S4–S8. Analyses were conducted using the JTT model of amino acid substitution^47^ with a gamma distribution (shape parameter = 1) of among-site rate variation. This analysis utilized 26 amino acid sequences. All ambiguous amino acid positions were removed for each sequence pair (i.e., pairwise deletion option). The final data set contained 8,105 amino acid positions in ORF1ab, 1,736 positions in S, and 94 positions in E. All evolutionary analyses were conducted in MEGA11^48,49^.

### Phylogenetic analysis

To reveal the phylogenetic relationships among betacoronaviruses, representative virus genomes were obtained from NCBI/GenBank (https://www.ncbi.nlm.nih.gov/). Identical sequences were removed using CD-HIT-EST version 4.8.1^50^ and the remaining nucleotide or amino acid sequences were aligned by using the L-INS-i program in MAFFT version 7.453^51^. Ambiguously aligned regions were trimmed using trimAl v1.5^52^, with a gap threshold of 0.9 and a minimum conservation threshold of 60%. Maximum likelihood phylogenetic trees were then inferred using IQ-TREE 2 v2.3.6^53^, with the optimal substitution model selected by ModelFinder^54^. Branch support was calculated using 1,000 bootstrap replicates^21^ with the UFBoot2 algorithm and an implementation of the SH-like approximate likelihood ratio test available within IQ-TREE 2.

### Prediction of furin cleavage sites near the S1/S2 junction in betacoronavirus S proteins

In June 2025, we downloaded all full-length protein sequences annotated as “*Betacoronavirus*” or “unclassified *Betacoronavirus*” from GenBank. To avoid redundancy, all SARS-CoV-2 sequences were excluded with the exception of one representative sequence. We then extracted sequences whose protein descriptions contained the keywords “S protein,” “spike,” or “surface glycoprotein,” yielding a total of 1,923 sequences. Of these, 1,622 sequences with annotated host information were retained. Redundant sequences were removed using CD-HIT-EST version 4.8.1^50^, and the BRZ batCoV sequence was manually added, resulting in a final data set of 935 sequences for analysis. To predict potential furin cleavage sites (FCSs) near the S1/S2 junction of the spike protein, we analyzed the amino acid region spanning residues 599– 900 using ProP v1.0b (ProPeptide Cleavage Site Prediction)^55^. This region was chosen to cover the putative S1/S2 junction where furin-mediated cleavage typically occurs in betacoronaviruses. A site was considered a putative FCS if the ProP score exceeded the default threshold.

### Protein structure prediction

Spike protein structures for BRZ batCoV and select betacoronaviruses were predicted using the AlphaFold3 web server using default settings^22^. Five predictions for each model were considered and, in all cases, the top ranked prediction by AlphaFold3 based on overall structural confidence scores was selected. Structural homology across spike proteins was evaluated using structural superpositions performed using the FATCAT webserver (version 2.0)^56^ and the Matchmaker tools using the Needleman-Walsh alignment algorithm^57^ and best chain pairing within UCSF ChimeraX (version 1.10)^58^. Protein structures were visualized and annotated by using UCSF ChimeraX.

### Cells

Expi293F cells (Thermo Fisher Scientific) were maintained in the HE400AZ medium (Gmep Inc.) at 37°C in 8% CO_2_. The cells were tested for mycoplasma contamination using PCR and were confirmed to be mycoplasma free.

### Plasmids

The codon-optimized spike genes of the BRZ batCoV, SARS-CoV-2 Wuhan-Hu-1 (GenBank: QHD43416), and bat coronavirus RaTG13 (GenBank: QHR63300.2) were designed for expression in mammalian cells and synthesized from GeneArt DNA Synthesis (Thermo Fisher Scientific). DNA sequences encoding the spike ectodomain of BRZ batCoV (amino acid residues 1–1418), SARS-CoV-2 (amino acid residues 1–1208), and RaTG13 (amino acid residues 1–1204), which include a C-terminal foldon trimerization motif followed by an octa-histidine tag, were cloned into the pcDNA3.1(+) expression vector (Invitrogen) and designated as pcDNA3.1-BRZ.batCoV-S-Foldon-8×His, pcDNA3.1-SARS-CoV-2-S-Foldon-8×His, and pcDNA3.1-RaTG13-S-Foldon-8×His, respectively.

Mutant spikes with a modified FCS were generated by substituting the FCS motif in BRZ batCoV (RDAR at amino acid residues 828–831) and SARS-CoV-2 spike (RRAR at amino acid residues 682–685) with the motif “GSAS” by using site-directed mutagenesis. A double proline (PP) mutation (T1179P and L1180P for BRZ batCoV; K986P and V987P for SARS-CoV-2) was also introduced to stabilize the pre-fusion conformation of the trimeric spike protein^59^. All constructs were confirmed by DNA sequencing.

### Preparation of purified spike proteins

Expi293F cells were transiently transfected with plasmids encoding recombinant spike proteins - (i) the wild-type, the GSAS mutant, and the GSAS/PP mutant BRZ batCoV spikes, (ii) the wild-type and the GSAS/PP mutant SARS-CoV-2 spike, and (iii) the wild-type RaTG13 spike - by using the Gxpress 293 Transfection Kit (Gmep Inc.) according to the manufacturer’s instructions. Five days post-transfection, cell culture supernatants were collected, clarified using a 0.45-µm syringe filter, and subjected to Ni-NTA affinity chromatography (Ni Sepharose 6 Fast Flow, Cytiva). After being washed with a buffer containing 50 mM sodium phosphate, 500 mM NaCl, and 40 mM imidazole at pH 7.4, the His-tagged proteins were eluted with a buffer containing 50 mM sodium phosphate, 150 mM NaCl, and 300 mM imidazole at pH 7.4. The eluates were subsequently concentrated using VivaSpin20, 100K MWCO (Sartorius) and subjected to size-exclusion chromatography using a Superdex 200 Increase 10/300 GL column (Cytiva) equilibrated with a 50 mM HEPES buffer containing 150 mM NaCl at pH 7.4. The concentration of the spike proteins was spectroscopically determined by absorbance measurements at 280 nm using 183480, 135845, and 137335 M^-1^ cm^-1^ for BRZ batCoV, SARS- CoV-2, and RaTG13, respectively, as a molar extinction coefficient. Molar extinction coefficients were predicted using ProtParam (https://web.expasy.org/protparam)^60^. The purified spike proteins were treated with PNGase F (NEB) by following the manufacturer’s protocol under non-denaturing and denaturing conditions.

### Electrophoresis

SDS-PAGE was conducted using a 5%–20% precast polyacrylamide gel (e-PAGEL E-T520L, ATTO). Samples were mixed with 5% (*v*/*v*) 2-mercaptoethanol and boiled at 100°C for 10 minutes prior to electrophoresis. The gels were then stained with Coomassie brilliant blue R- 250 (Wako) or EzStain silver (ATTO). For western blot analysis, after SDS-PAGE, the separated proteins in the gel were blotted onto a 0.45-µm PVDF membrane (Immobilon-P, Millipore) and blocked with 5% (*w*/*v*) skim milk in phosphate-buffered saline containing 0.1% Tween20 (PBS-T). The spike proteins were detected using a mouse monoclonal anti-His tag antibody (1:5000; D291-3, MBL) as the primary antibody and an HRP-conjugated anti-mouse IgG antibody (1:5000; Jackson) as the secondary antibody. The blots were developed with ImmunoStar Zeta chemiluminescent reagent (Wako).

## Acknowledgments

We thank Susan Watson for scientific editing, Mikiko Tanaka, Tomomi Kirino and Yurie Kida for technical assistance, and Kanako Hiromatsu for support with grant procedures at the University of Osaka. We thank So Nakagawa and Shintaro Shichinohe for discussions regarding data analysis and/or RNA pooling. We thank CAPES (Coordenação de Aperfeiçoamento de Pessoal de Nível Superior) for support. This study was also supported by KAKENHI Grants-in-Aid for Scientific Research on Innovative Areas [16H06429 (to T.W.), 16K21723 (to T.W.) and 16H06434 (to T.W)]; a Grant-in-Aid for Scientific Research (B) [JP22H02521 (to T.W.)]; Grant-in-Aid for Early-Career Scientists [25K18814 (to K.T.) and 22K15469 (to K.T.)]; Grant-in-Aid for JSPS Fellows 21J01036 (to K.T.); the AMED Research Program on Emerging and Re-emerging Infectious Diseases [JP19fk0108113 (to T.W.) and JP19fk018113 (to T.W.)]; AMED under Grant Numbers JP223fa627002, JP22am0401030, and JP23fk0108659 (to T.W.); AMED Advanced Research and Development Programs for Medical Innovation (AMED-CREST) 22gm1610010h0001 (to T.W.); the Takeda Science Foundation (to T.W.); RIKAKEN HOLDINGS CO. Young Researcher Support Grant-in-aid (to K.T.); and a National Health and Medical Research Council (Australia) Investigator grant (GNT2017197) to E.C.H.

## Contributions

K.T. and T.W. conceptualized this study. T.M., F.B.C., E.T., M.M., T.JS. L., J.M and T.W. performed the animal sampling and processed the samples. K.T. analyzed sequence data. N.Y. and I.A. performed viral protein expression experiments. K.T. and J.C.O.M conducted protein structural modeling analysis. K.T., N.Y, J.C.O.M., E.C.H., and T.W. interpreted the data. K.T. wrote the original draft. N.Y., J.C.O.M., E.C.H. and T.W. contributed to manuscript revisions and additions. K.T., Y.K. and T.W. organized international collaboration. All authors reviewed and approved the final manuscript.

## Competing interests

Y.K. has received unrelated grant support from Daiichi Sankyo Co., Ltd., Fujifilm Toyama Chemical Co., Ltd., Tauns Laboratories, Inc., Shionogi & Co. Ltd., Otsuka Pharmaceutical Co., Ltd., KM Biologics Co. Ltd., Kyoritsu Seiyaku Corporation, Shinya Corporation and Fuji Rebio, Inc. Y.K. is a co-founder of FluGen. The other authors do not have any competing interests.

## Data Availability Statements

The raw sequence reads generated for this project have been deposited at the Sequence Read Archive (SRA) database under Bioproject: PRJNA1345860 (BioSample accessions SAMN52820448 – SAMN52820469) and GenBank database (accession numbers: XXXX).

